# Construction of a Genetic Sexing Strain for *Aedes albopictus*: a promising tool for the development of sterilizing insect control strategies targeting the tiger mosquito

**DOI:** 10.1101/274712

**Authors:** Cyrille Lebon, Aude Benlali, Célestine Atyame, Patrick Mavingui, Pablo Tortosa

**Author notes:** Equal contribution (Cyrille Lebon, Aude Benlali). **Corresponding author:** UMR PIMIT, Université de La Réunion, INSERM 1187, CNRS 9192, IRD 249, Plateforme de Recherche CYROI, 2 rue Maxime Rivière, 97490 Ste Clotilde, La Réunion, France.

## Abstract

**Background:** *Aedes albopictus* is an invasive mosquito species of global medical concern as its distribution has recently expanded to Africa, the Americas and Europe. In the absence of prophylaxis protecting human populations from emerging arboviruses transmitted by this mosquito species, the most straightforward control measures rely on the suppression or manipulation of vector natural populations. A number of environmental-friendly methods using innundative releases of sterilizing males are currently under development. However, these strategies are still lacking an efficient sexing method required for mass production of males.

**Results:** We present the first Genetic Sexing Strain (GSS) in *Ae. albopictus*, hereafter referred as TiCoq, obtained by sex linkage of *rdl* gene conferring dieldrin resistance. Hatching rate, larval survival and sex ratio were followed during twelve generations. The use of dieldrin at third larval stage allowed selecting 98% of males on average.

**Conclusion:** A good production rate of TiCoq males makes this GSS suitable for any control method based on mass production of *Ae. albopictus* sterilizing males. Despite limitations resulting from affected egg hatch as well as the nature of the used insecticide, the construction of this GSS paves the way for industrial sex separation of *Ae. albopictus*.

## Introduction

The recent Zika virus (ZIKV) epidemic has put vector borne diseases under the spotlight, and further confirmed the growing concern of emerging vector-borne pathogens [1]. Somehow, this epidemic shows similarities with Chikungunya virus (CHIKV) emergence a decade ago: both CHIKV (Alphavirus) and ZIKV (Flavivirus) were initially known to cause outbreaks limited in space and time before firing at a global scale. These epidemics tend to remind that we will most certainly experience future epidemics caused by arboviruses mostly unknown to the scientific community. As a consequence, the prophylactic treatments needed for the control of such diseases will be either scarce or absent.

One way to prevent the spread of emerging arthropod-borne pathogens is to limit their transmission through the control of their invertebrate vectors. Such control methods, which have been so far mostly implemented through the use of synthetic pesticides, have proven efficient and notably allowed eradicating malaria from several territories worldwide following World War II. However, the recurrent selection of insecticide resistance in vector natural populations [2], together with unwanted effects on non-target species has conducted restriction uses of synthetic pesticides and stimulated the development of a handful of environmentally friendly methods. The field of innovative vector control development is currently exceptionally dynamic [3], and several ongoing programs have reached the step of field pilot trial assays [4,5]. Among the strategies aiming at suppressing mosquito natural populations, sterilizing technologies based on the innundative release of sterilizing males have been experiencing a spectacular revival. These technologies use either Gamma- or X- ray irradiation [6], transgenesis [7], *Wolbachia* symbiotic bacteria [8] or more sophisticated associations such as the recently proposed boost Sterile Insect Technique, in which males are sterilized through irradiation and then used as dispersers of entomopathogenic viruses and/or insecticides [9]. All these technologies rely on the availability of an efficient sexing method allowing the separation of males and females at industrial scales, which is compulsory for the innundative release of treated males.

The control of vector populations through the release of sterile males has proven successful for crop pest species such as *Cochliomyia hominivorax* or *Ceratitis capitata* [10]. By contrast, these vector control strategies have not been implemented at industrial scales for the control of mosquitoes. The availability of an efficient sexing method is indeed one of the main technological locks. Thus far, pilot trials have been using manual or mechanical separation techniques, which are not either up scalable or fully satisfactory [11]. Indeed, mechanical separation relies on sexual dimorphism, which can allow efficient sieving as reported for *Ae. aegypti* [7]. However, sexual dimorphism may not be sharp enough in some species, such as *Ae. albopictus*, for which sieving is notoriously challenging and results in important losses of produced male pupae [12,13].

Among the several methods currently explored for sex separation in *Ae. albopictus* [14], we used a dieldrin conferring marker to construct a Genetic Sexing Strain (GSS), essentially adapting a strategy previously used for *Culex tarsalis* [15] and *Anopheles arabiensis* [16]. Sex linkage of the *rdl* gene, conferring dieldrin resistance, was obtained through X-ray irradiation and selection of the sex biased mosquito lines. The data presented herein provide, to our knowledge, the first GSS reported to date for *Ae. albopictus*, and hereafter referred as TiCoq. We present the performance and some life history traits of TiCoq and discuss future improvements that will accelerate the implementation of sterilizing strategies targeting the tiger mosquito.

## Materials and methods

### Mosquito stocks and rearing

Two *Ae. albopictus* lines originally sampled in two localities on La Réunion Island (*Plaine des Palmistes* and *La Providence*) were selected for this study. These localities correspond to an elevated rural area and a coastal urban park of La Réunion Island sheltering natural mosquito populations displaying low and high *rdl*^*R*^ allelic frequencies, respectively [17]. Larvae were reared at a density of approximately 1000 first instar larvae in white plastic trays (30×40 cm with a depth of 6 cm) containing 3L of distilled water and were fed ad libitum with a mixture of rabbit and fish-food (Tetra GmbH, TetraMin, Heinsberg, Germany). Upon pupation, males and females were separated manually under a binocular loop using terminalia dimorphism and placed into insect rearing cages (30×30×30 cm) until emergence. Adults were fed with 10% sucrose solution [w/v] and females were blood-fed using the Hemotek feeding system (Discovery Workshops, Lancashire, United Kingdom) and defibrinated cow blood. The use of blood provided by the slaughterhouse for mosquito feeding does not require ethical clearance.

### Bioassays

A 1,000 ppm dieldrin (Dr Ehrenstorfer, Germany) stock solution was prepared in absolute ethanol and used as a stock for further dilutions. Batches of 25 third instar larvae were placed in 100 ml of water containing 0 to 10 ppm dieldrin. Three replicates were prepared for each insecticide concentration and cups were conserved at room temperature. All surviving and dead larvae were counted in each cup after 24 hours and data were analysed using BioRssay 6.2 [18].

### Molecular typing

Adult mosquito DNA was extracted as previously described [19]. A PCR-RFLP test [17] was used to detect the Ala to Ser substitution in *rdl* conferring dieldrin resistance in *Cx. pipiens* and *Ae. albopictus*. PCR was run for 30 cycles (94°C for 30 s, 52°C for 30 s, and 72°C for 1 min), followed by 5 min at 72°C. The PCR product was then digested for 3 hours at 60°C with *BstAPI* restriction endonuclease (New England Biolabs, Evry, France), which selectively cleaves the susceptible allele [17]. Allelic profiles were then visualized using 2% agarose electrophoresis of digested PCR product stained with Gelred.

### X-Ray irradiation and GSS identification

Chromosomal translocations were induced using a BloodXrad X-ray irradiator (Cegelec, Bretigny sur Orge, France) located at the Institut Français du Sang, Centre Hospitalier Universitaire de La Réunion. For this, 100 *rdl* homozygous resistant (R-Run) male pupae were placed in a jar (Ø: 11cm; H: 8.5cm) containing 40 ml distilled water and were irradiated with one of the two selected irradiation doses: 20Gy and 25Gy. Each of the two batches of males, corresponding to the two irradiation doses, were then crossed *en masse* with *rdl* homozygous susceptible (S-Run) females, leading to the F0 heterozygous generation. Forty crosses were performed using one F0 (heterozygous) male and five S- Run females into a small insect holding cage (12×12×12 cm). Eggs from each cage were allowed to hatch and up to 50 third instar larvae were placed at room temperature for 24h in plastic cups containing 100 mL of water supplemented with 0.1 ppm dieldrin. Sex ratio was determined at pupal stage and all broths displaying >60% males following dieldrin selection were allowed to emerge. Males from each progeny were then backcrossed with S-Run females and dieldrin pressure was maintained at each following backcross.

## Results

### Construction of dieldrin resistant (R-RUN) and susceptible (S-RUN) homozygous mosquito lines

The construction of a GSS through sex linkage of a selectable insecticide resistance gene requires the availability of homozygous resistant and susceptible mosquito lines. A dieldrin sensitive strain was constructed as follows: 20 cages were set with five females and one male in each cage. Adult mosquitoes were left for 48 hours in the presence of cotton soaked in 10% sucrose. Each male was then genotyped at the *rdl* locus following a previously described procedure [17]. Only those cages whith homozygous *rdl*^*S*^ males were conserved and females were given a blood meal. Two days later, each female was placed in a separate cage and allowed to lay eggs, which were conserved while females were genoytyped. Those eggs laid by *rdl*^*S*^ homozygous females were poolled and immersed in water for hatching, providing an homozygous sensitive line hereafter named S-Run.

The construction of an homozygous resistant strain required two rounds of dieldrin selection to increase the frequency of *rdl*^*R*^ allele in the cages. For this, third and fourth instar larvae from *La Providence* were placed in plastic cups containing 50 mL of water with increasing dieldrin concentration. F0 larvae were placed in 0.05 ppm dieldrin for 24h and surviving larvae were subsequently fed until emergence. Eggs laid by surviving F0 females were allowed to hatch and third and fourth instar larvae were placed in plastic cups filled with 50 mL of water containing 0.1 ppm dieldrin. Surviving F1 larvae were then allowed to moult and nymphs were sex separated. Isofemale lines were constructed following the same exact procedure as that implemented for the construction of S-Run, and eggs resulting from copulations between homozygous *rdl*^*R*^ male and female mosquitoes were pooled. Hatching eggs led to the homozygous *rdl*^*R*^ mosquito line hereafter referred as R-Run.

### Determination of the dieldrin diagnostic dose

In addition to pure-breeding susceptible and resistant homozygous larvae, heterozygous F1 [RS] larvae were obtained by crossing R-Run males with S-Run females. Batches of 25 third instar larvae of homozygous susceptible [SS], heterozygous [RS] or homozygous resistant [RR] mosquitoes were placed in 100 mL of water containing 0 to 10 ppm dieldrin, and left for 24h at room temperature. Surviving and dead larvae were counted for each insecticide dose (3 cups for each genotype/dieldrin dose). Survival was plotted for each dieldrin dose and mosquito genotype. As shown on figure 1, a dieldrin concentration of 0.1 ppm killed 100% of S-Run larvae but none of the heterozygous or R-Run larvae, and was thus selected as the diagnostic/selective dose.

**Figure 1:**
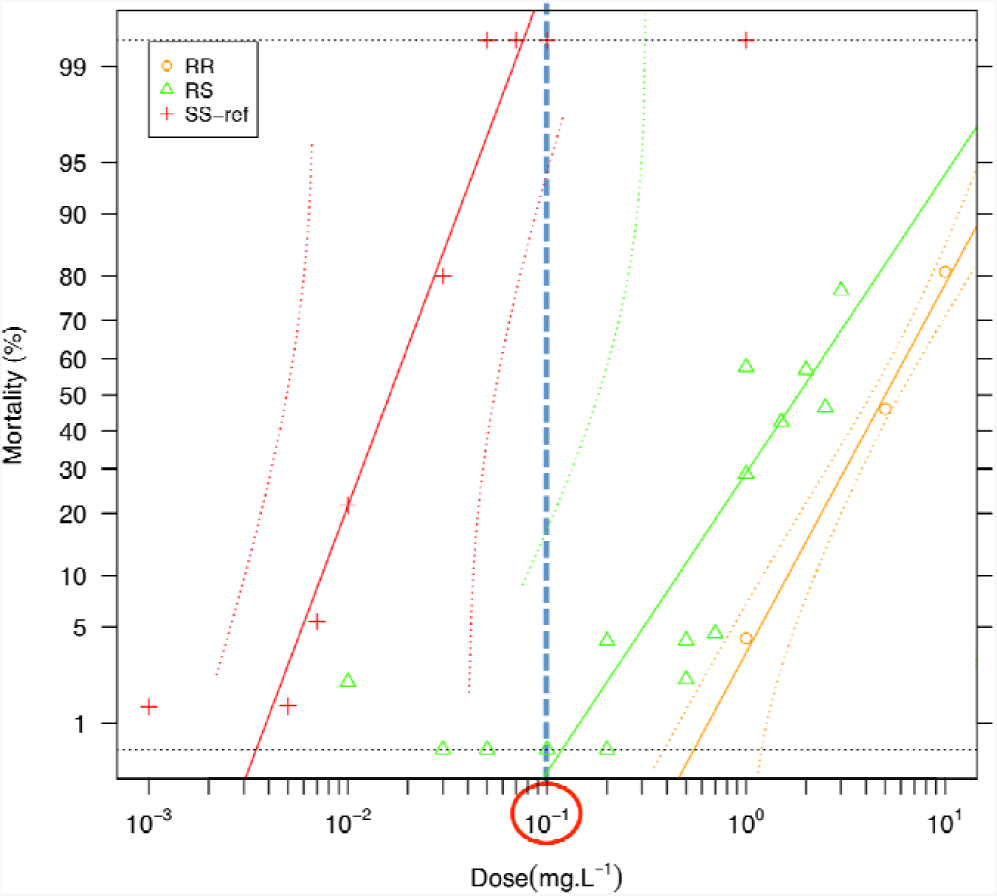
Survival of R-Run (RR), S-Run (SS) and heterozygous (RS) third instar larvae following 24h treatment with increasing dieldrin concentrations. The diagnostic dose is circled.

### X-Ray irradiation, identification and amplification of TiCoq

Following irradiation at 20 or 25Gy, R-Run male pupae were allowed to emerge and then crossed *en masse* with virgin S-Run females for 48h, females were subsequently blood fed and allowed to lay eggs. As expected, hatching rates were significantly lower in crosses involving males irradiated at 25Gy than in crosses involving males irradiated at 20Gy (93.38 ± 0.99 and 82.89 ± 3.33, respectively; Student test: t=3.53 P=0.001). Following dieldrin selection, no difference was observed between these two irradiation doses either for larvae survival rate (20Gy: 47.87 ± 1.19; 25Gy: 49.61 ± 1.49) or sex ratio (20Gy: 50.48 ± 1.52; 25Gy: 54.70 ± 2.29). Forty F1 dieldrin resistant males (20 males for each irradiation dose) were then individually crossed with S-Run females (one male and five females per cage), blood fed and allowed to lay eggs. Only four males led to progenies exhibiting male biases exceeding 60% and were further backcrossed. As shown on Table 1, a single male produced strong male bias (100%) over 4 successive backcrosses. This mosquito line, hereafter referred as TiCoq, was maintained and amplified while all other lines were discarded. At each generation, TiCoq males were back-crossed with S-Run females with a 1:3 male/female ratio. Hatching and survival rates as well as sex ratio were measured for each generation. As shown on figure 2, hatching rate averaged 30% over the entire experiment, larvae surviving rate following dieldrin treatment averaged 50% and sex ratio bias was stable at 97.8% (SD 5.96).

**Table 1:** *Aedes albopictus*, dieldrin selection of Genetics Sexing Strain candidates.

| F0 Irradiation dose (Gy) | Generation | Starter male | Eggs number | Hatching rate (%) | Number of larvae exposed to dieldrin | Survival rate following dieldrin selection (%) | Number of male pupae | Number of pupae | Male % |
| --- | --- | --- | --- | --- | --- | --- | --- | --- | --- |
| 20 | F2 | A | 169 | 95.27 | 138 | 48.55 | 40 | 66 | 60.61 |
| 20 | F2 | B | 139 | 93.53 | 122 | 51.64 | 37 | 56 | 66.07 |
| 25 | F2 | C | 143 | 90.21 | 114 | 41.23 | 24 | 33 | 72.73 |
| 25 | F2 | D | 121 | 30.58 | 18 | 44.44 | 6 | 6 | 100.00 |
| 20 | F3 | A | 192 | 88.02 | 117 | 62.39 | 44 | 70 | 62.86 |
| 20 | F3 | B | 352 | 86.93 | 251 | 51.79 | 64 | 116 | 55.17 |
| 25 | F3 | C | 486 | 74.07 | 316 | 47.78 | 68 | 139 | 48.92 |
| 25 | F3 | D | 149 | 24.16 | 31 | 29.03 | 7 | 7 | 100.00 |
| 20 | F4 | A | 1242 | 25.93 | 284 | 50.00 | 62 | 141 | 43.97 |
| 25 | F4 | D | 929 | 13.46 | 104 | 46.15 | 36 | 36 | 100.00 |
| 25 | F5 | D | 1465 | 34.61 | 390 | 47.95 | 125 | 132 | 94.70 |
| 25 | F6 | D | 6462 | 28.04 | 1540 | 47.14 | 500 | 509 | 98.23 |
| 25 | F7 | D | 2288 | 31.42 | 538 | 37.55 | 80 | 82 | 97.56 |
| 25 | F8 | D | 482 | 26.56 | 95 | 37.89 | 30 | 30 | 100.00 |
| 25 | F9_a | D | 1218 | 32.02 | 317 | 48.90 | 102 | 115 | 88.70 |
| 25 | F9_b | D | 1747 | 27.48 | 429 | 48.25 | 113 | 116 | 97.41 |
| 25 | F10 | D | 1499 | 34.89 | 350 | 49.71 | 164 | 166 | 98.80 |
| 25 | F11 | D | 3241 | 39.46 | 1648 | 46.84 | 744 | 750 | 99.20 |
| 25 | F12 | D | 2461 | 35.19 | 759 | 47.56 | 340 | 345 | 98.55 |

**Figure 2:**
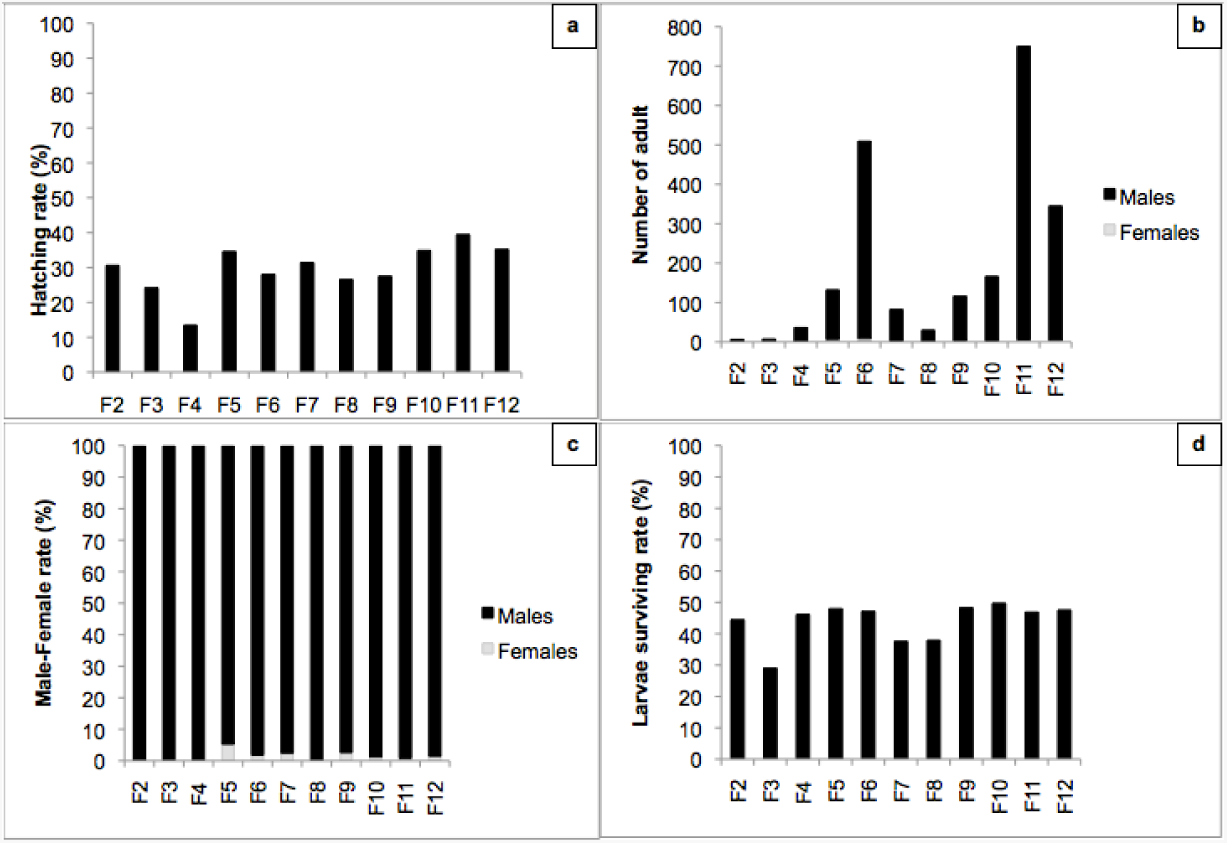
TiCoq life history traits. Parameters collected over 12 generations under dieldrin pressure. a: Hatching rate; b: number of surviving adults; c: sex ratio and d: larvae surviving rate.

### Fecundity of TiCoq females obtained following insecticide pressure

Sex bias was not perfect and 2% females descending from crosses between TiCoq males and S-Run females survived the dieldrin treatment at larval stage. Seven out of these surviving females were genotyped showing that they were all heterozygous at *rdl* (Table 2). This pattern suggests that these females actually resulted from meiotic recombination (see discussion). When these females were crossed with sensitive S-Run males, hatching rates were high (84,82% ± 6,91), sex ratio was not biased as dieldrin treatment of third instar larvae led to 50,33 % (± 18,12) males (see Table 2).

**Table 2:** Life history traits of larvae resulting from TiCoq dieldrin resistant females crossed with S-Run males.

| Generation | Female ID | Hatching rate (Total number of eggs) | Genotype | Number of larvae exposed to dieldrin | Survival rate following dieldrin selection (%) | Number of male pupae | Number of female pupae | Male % |
| --- | --- | --- | --- | --- | --- | --- | --- | --- |
| F10 | A | 100,00 (4) | RS | NT | NT | NT | NT | NT |
| F10 | B | 98,90 (91) | RS | NT | NT | NT | NT | NT |
| F10 | C | 92,86 (14) | RS | NT | NT | NT | NT | NT |
| F11 | D | 48,28 (29) | RS | 9 | 22.22 | 2 | 0 | 100.00 |
| F11 | E | 74,31 (109) | RS | 58 | 48.28 | 14 | 14 | 50.00 |
| F11 | F | 92,96 (71) | RS | 39 | 38.46 | 2 | 12 | 14.29 |
| F11 | G | 86,44 (59) | RS | 52 | 53.85 | 10 | 17 | 37.04 |

### Stability of TiCoq

In order to test a possible drift of TiCoq in the absence of selective pressure, a part of layers was not treated with dieldrin after the 7^th^ generation. For this, males and females were kept in rearing cages following emergence and the line was maintained without insecticide pressure during all subsequent generations while 50-100 3rd or 4^th^ instar larvae were randomly sampled and treated with dieldrin at each generation in order to measure *rdl*^*R*^ frequency. Egg hatch steadily increased from 25.7% to 92.7% over 4 generations while larvae survival following dieldrin treatments decreased from 47.14% to 3.20% in the meantime. No resistant larvae were observed after 5 generations in the absence of insecticide pressure (Table 3).

**Table 3:** TiCoq performance without dieldrin selection.

| Generation | Eggs number | Hatching rate (%) | Number of larvae exposed to dieldrin | Survival rate following dieldrin selection (%) | Number of male pupae | Number of pupae | Male % |
| --- | --- | --- | --- | --- | --- | --- | --- |
| F1 | 1219 | 25.68 | 280 | 47.14 | 70 | 70 | 100.00 |
| F2 | 2147 | 40.06 | 627 | 29.98 | 159 | 160 | 99.38 |
| F3 | 1373 | 64.02 | 850 | 8.35 | 64 | 65 | 98.46 |
| F4 | 328 | 92.68 | 250 | 3.20 | 7 | 7 | 100.00 |
| F5 | 1292 | 93.73 | 1102 | 0.00 | / | / | / |

## Discussion

We report the first GSS for *Ae. albopictus*. TiCoq allows producing 97.8% males following dieldrin treatment of third instar larvae. Such a genetic tool may advantageously replace traditional sieving that has proven imperfect for this species [12,13]. TiCoq hence paves the way for the up scaling of incompatible/sterile insect techniques targeting *Ae. albopictus*. The measurement of some life history traits shows that the line can be easily maintained in the presence of dieldrin selection. However, TiCoq still displays a number of drawbacks that need to be discussed and for which improvements must be considered.

The strong bias in sex ratio provided by TiCoq could be considered as strong enough for traditional sterile insect technique, although proper modelling will need to be carried out at each future environmental set-up chosen for pilot trials. By contrast, incompatible insect technique [20] requires virtually no survival of females following treatment. Indeed, released females will be fertile with incompatible males, and may lead to the invasion of an hetereospecific *Wolbachia* in *Ae. albopictus* natural populations [5]. As shown by the characterization of surviving TiCoq females following dieldrin selection, all tested females were heterozygous for *rdl*. In addition, these females were nearly fully fertile when crossed with non-translocated S-Run males. The recovery of a nearly perfect fecundity suggests that these females result from a meiotic recombination restoring chromosome integrity rather than a recombination between the male locus and the linked *rdl*. Anyway, a good fitness of these RS TiCoq females will undoubtedly provide any accidentally released females with good chances of laying viable eggs. As a consequence, R-Run translocation will have to be repeated as long as the obtained sex ratio distortion is not close to 100%.

The main drawback of this GSS is certainly the used marker, as dieldrin has been banned for decades in a number of countries. It must be emphasized that in no circumstances the insecticide should be used in the field. Larvae or even eggs [21] should be treated exclusively in the laboratory for sexing purposes. However, traces of dieldrin may persist in adults as previously reported in *Anopheles arabiensis* GSS [22]. This could be circumvented by different ways. First, dieldrin could be replaced by other pesticides, such as fipronil, for which cross-resistance with Gaba receptor mutants has been previously reported [23–25]. Second, the present work shows the feasibility of genetic translocation for the generation of a sex separation tool in *Ae. albopictus*. The mutation conferring dieldrin resistance is thus far one of the few available robust selectable marker for this mosquito species. However, given the medical importance of this mosquito species, it is likely that other insecticide resistance markers (such as the recently reported *kdr* mutants [26] or carboxylesterase gene amplifications [27]) will be fully characterized in the next future and will be made available for the development of a GSS avoiding the use of banned insecticides. Lastly, sterile insect strategies are appealing as they are both highly specific and environmentally friendly. Therefore, a high throughput screen allowing the identification of “clean” markers may be a profitable investment. Temperature sensitive lethal mutations have been previously discovered for other Culicinae species [28] and used at industrial scales for the control of medflies [29]. Hence, the present work shows that the identification or the development of such a marker by random mutagenesis would pave the way to the implementation of fully clean sterile insect strategies that are urgently needed for the control of future emerging epidemics involving the tiger mosquito.

## Acknowledgements

We wish to thank Hafsah Limbada, Emmanuelle Clervil and Mathilde Pagès for their help in the construction of R-Run and S-Run mosquito lines. Institut Français du Sang is warmly acknowledged for giving permanent access to the irradiator facilities. We thank Jean-Sébastien Dehecq and Abdoul Rutee for providing larvae from Plaine des Palmistes that allowed the establishment of the S-Run mosquito line. This work was supported by a Credit Impôt Recherche provided by the French Ministry of research and higher education to SymbioTIC company, and by the joint FAO/IAEA Coordinated Research Program “Exploring genetic, molecular, mechanical and behavioural methods of sex separation in mosquitoes” (http://www-naweb.iaea.org/nafa/ipc/crp/ipc-behavioural-methods.html).

